# Sparse Testing Designs for Optimizing Predictive Ability in Sugarcane Populations

**DOI:** 10.1101/2024.03.14.584687

**Authors:** Julian Garcia-Abadillo, Paul Adunola, Fernando S. Aguilar, Jhon Henry Trujillo-Montenegro, John Jaime Riascos, Reyna Persa, Julio Isidro y Sanchez, Diego Jarquín

**Affiliations:** Centro de Biotecnología y Genómica de Plantas, Universidad Politécnica de Madrid, Madrid, Spain; Agronomy department, University of Florida, Gainesville, FL, USA; Horticultural Sciences Department, University of Florida, Gainesville, FL 32611; Centro de Investigación de la Caña de Azúcar de Colombia, Florida-Cali, Colombia

**Keywords:** Genomic Selection GS, Genomic Prediction GP, Sparse Testing Designs, Optimization, Sugarcane breeding

## Abstract

Sugarcane is a crucial crop for sugar and bioenergy production. Saccharose content and total weight are the two main key commercial traits that compose sugarcane’s yield. These traits are under complex genetic control and their response patterns are influenced by the genotype-by-environment (G×E) interaction. An efficient breeding of sugarcane demands an accurate assessment of the genotype stability through multi-environment trials (METs), where genotypes are tested/evaluated across different environments. However, phenotyping all genotype-in-environment combinations is often impractical due to cost and limited availability of propagation-materials. This study introduces the sparse testing designs as a viable alternative, leveraging genomic information to predict unobserved combinations through genomic prediction models. This approach was applied to a dataset comprising 186 genotypes across six environments (6 × 186 = 1,116 phenotypes). Our study employed three predictive models, including environment and genotype as main effects, as well as the G×E interaction to predict saccharose accumulation (SA) and tons of cane per hectare (TCH). Calibration sets sizes varying between 72 (6.5%) to 186 (16.7%) of the total number of phenotypes were composed to predict the remaining 930 (83.3%). Additionally, we explored the optimal number of common genotypes across environments for G×E pattern prediction. Results demonstrate that maximum accuracy for SA (*ρ* = 0.611) and for TCH (*ρ* = 0.341) was achieved using in training sets few (3) to no common (0) genotype across environments maximizing the number of different genotypes that were tested only once. Significantly, we show that reducing phenotypic records for model calibration has minimal impact on predictive ability, with sets of 12 non-overlapped genotypes per environment (72 = 12 × 6) being the most convenient cost-benefit combination.

## 1 Introduction

The world faced an urgent challenge to provide nutritional sustenance for a burgeoning 8 billion people (UNPF, 2023) amidst threats to food security, climate change, and finite resources (UNCCD, 2017; FAO, 2015, 2018). Bridging the gap between food production and population growth necessitates innovative agricultural solutions. Sugarcane emerges as a key player in this scenario, serving dual purpose: *i*) as a primary sugar source (a dietary staple), and *ii*) bioenergy feedstock production (Goldemberg, 2008; Waclawovsky et al., 2010). The versatility of sugarcane makes it an ideal crop for targeted breeding efforts to enhance both yield (sugar and biofuel) (Hoang et al., 2015) and yield stability (Scortecci et al., 2012). However, sugarcane breeding is challenged by long breeding cycles, low genetic diversity, large genome size and clonal propagation, limiting efficient genetic improvement and yield potential (Souza et al., 2011; Hoang et al., 2015; Yadav et al., 2020).

In plant breeding, identifying superior cultivars for agricultural demands remain critical. Sugarcane’s narrow genetic base, primarily due to vegetative propagation, complicates breeding efforts (Roach, 1989; Raboin et al., 2008; Wei et al., 2016). Multi-environmental trials (METs) are essential in sugarcane breeding, enabling the evaluation of genotype performance across environments and identifying stable or specifically adapted cultivars (Jackson et al., 1992; Abu-Ellail et al., 2020). However, METs are resource-intensive, particularly for clonally propagated crops, prompting the need for more efficient trial designs.

Genomic selection (GS) has been transformative in plant breeding, enhancing genetic gain rates over the past two decades (Crossa et al., 2010, 2011, Jarquin et al., 2014; Ferrão et al., 2021). GS uses genome-wide molecular markers in genomic prediction (GP) models to estimate breeding values of untested individuals (Meuwissen et al., 2001). By capturing the genetic variation in the genome, GP enables the prediction of complex traits that are otherwise challenging to assess using conventional breeding tools (Yadav et al., 2020; Ferrão et al., 2021). This approach facilitates early-stage breeding cycle decisions, reducing the need for extensive field trials and expediting cultivar development.

Genomic prediction is particularly advantageous in predicting performance of individuals in new and partially tested environments through cross-validation (CV) schemes. This practice ensures that the predictive ability of GS models extends beyond the environments in which genotypes are bred, allowing accurate selection of individuals with potential high performance across a range of environmental and management conditions. Empirical studies suggest that incorporating G×E interaction in GP models can streamline the breeding process, either by skipping certain stages or by reducing the number of genotypes tested in fields, thereby enhancing trial testing capacity (Crossa et al. 2017; Resende et al., 2018; Jarquin et al., 2020).

Sparse testing, the strategic selection of a subset of genotypes for evaluation in target environments, offers an innovative alternative to field trials for the entire population. This approach leverages statistical techniques and predictive models, such as GP, to extrapolate the performance of the entire population from a selectively observed subset. Through model calibration and cross-validation, sparse testing enables prediction of unobserved genotype-in-environment combinations. Consequently, it optimizes resource allocation, allowing breeders to focus on the most promising candidates from the target population of environments (TPEs). This methodology not only accelerates the breeding cycle but also the cost-effective identification of superior cultivars.

Sparse testing designs often mirror cross validation (CV) schemes CV1 and CV2 (Burgueno et al. 2012; Jarquin et al., 2017), encompassing a broad range of genotype-in-environment combinations. In the CV1 scenario, genotypes never tested at any environment are predicted, while CV2 involves predicting already observed genotypes in incomplete METs. This raises critical questions about the optimal design of sparse testing in METs, such as the balance between testing a few genotypes across multiple environments versus many genotypes in fewer environments; and the trade-off between prediction accuracy and selection intensity. Empirical evidence from crops like maize (Jarquin et al., 2020; Atanda et al., 2021; Montesinos-Lopez et al., 2023), wheat (Crespo-Herrera et al., 2021; Atanda et al., 2022) and soybean (Persa et al., 2023) has provided insights into optimizing sparse testing designs. Research by Jarquin et al. (2020) and Crespo-Herrera et al. (2021) revealed that GS model incorporating G×E interactions can maintain robust predictive ability even with reduced training sets. These studies indicate that the highest recovery of predictive ability in scenarios with either non-overlapping or completely overlapping genotypes in training set combinations mostly depends on the species.

In this context, this study investigates the predictive ability of different sparse-testing allocation designs in sugarcane breeding. We simulated the allocation of finite breeding resources to optimize the prediction of un-phenotyped genotypes in METs. For that, we varied the number of overlapping genotypes and training set sizes, similarly to the approaches of Jarquin et al. (2020) and Crespo-Herrera et al. (2021). We consider three allocation scenarios: *i*) Non-Overlapping Genotypes (NOG), with a single observation per genotype across environments, *ii*) Overlapping Genotypes (OG) where all training genotypes are phenotyped across all environments, resembling the CV1 scenario, and *iii*) a combination of the NOG and OG schemes. To assess the contribution of capturing G×E in METs, three models were fitted for each sparse testing case: *i*) main effect of environment and genotype (M1); *ii*) main effect of environment, genotype and genomic markers (M2); and *iii*) main effect of environment, genotype, genomic markers, and the interaction between genomic markers and environment (M3). Our findings demonstrate effective resource optimization strategies for METs in sugarcane breeding, potentially increasing testing capacity by fivefold within a fixed target set of genotype-in-environment combinations or reducing total phenotyping cost between 83% to 93%.

## 2 Materials and Methods

### 2.1 Plant Material and phenotyping

The dataset analyzed in this study comprises 220 genotypes (Jaimes et al., 2024) from the diverse panel of Centro de Investigación de la Caña de Azúcar, Cenicaña (Colombian Sugarcane Research Center). This panel includes 98 genotypes representing the phenotypic diversity of Cenicaña’s germplasm bank, 58 genotypes selected for their resistance or susceptibility to prevalent diseases and pests in Colombia, 33 genotypes encompassing introductions and commercial varieties, and 31 wild species from *Saccharum officinarum, S. spontaneum, S. barberi, S. sinensi,* and *Erianthus* spp. The genotypes were planted across three locations in the Valle del río Cauca, Colombia: Balsora (Mayaguez sugarcane mill) in 2016 (E1) and 2017 (E2), Taula Mejía (La Cabaña sugarcane mill) in 2018 (E3) and 2019 (E4), and Porvenir (Manuelita sugarcane mill) in 2020 (E5) and 2021 (E6). Each location was planted under an alpha-lattice design. Balsora and Taula Mejia were planted with 3 replications, while Porvenir, due to field limitations, had 2 replications. The experimental unit at each location comprised a plot of 5 rows, each 5 meters long and spaced 1.65 m apart. Commercial checks, genotypes S29, S177, and S64, were replicated multiple times within each replicate-block, resulting in a total of 717, 720, and 576 experimental units for Balsora, Taula Mejia, and Porvenir, respectively.

Phenotypic data for sucrose accumulation (SA) and tons of cane per hectare (TCH), were collected during harvest (13 months after planting). Sucrose accumulation was measured by shredding 10 stalks per experimental unit to obtain the juice per sample (Laharrahondo and Torres, 1989). Then the sucrose content was quantified from the extracted juice through a near infrared (NIR) spectrophotometer (Laharrahondo and Torres, 1989). For TCH, the measurement was taken after weighing all stalks per experimental unit. Both SA and TCH measurements were conducted for both plant cane and first ratoon at all locations.

### 2.2 Genotyping and quality control

DNA was extracted from each of the 220 genotypes following Dellaporta et al. (1983) protocol. Sequencing was conducted using both Genotyping-by-sequencing (GBS) and Restriction-site Associated DNA sequencing (RADSeq) on a Hiseq2000 Illumina System at a depth of 105X and 38X for GBS and RADSeq, respectively. Quality control was performed using FastQC (Andrews 2010) and Trimmomatic (Bolger et al. 2014). Subsequently, the cleaned data were aligned and mapped to the CC 01-1940 monoploid reference genome (Trujillo-Montenegro et al. 2021), utilizing Bowtie2-2.5 (Langmead and Salzberg 2012) with default parameters. SNP calling was conducted with NGSEP 4.0.2 (Tello et al. 2019) by filtering out markers that failed to meet the following quality control criteria: a Minor Allele Frequency (MAF) below 3%, Phred score below 30, distance between markers below 5 bp, presence of more than 50% of missing data, and an average depth below 5X, as well as markers with more than one allelic version. Post quality control, a finalized count of 22,324 SNP markers remained for analyses.

### 2.3 Phenotypic adjustment

The statistical model was independently applied to raw data at each environment aiming to adjust the genotype effect using the Best Linear Unbiased Predictor (BLUPs) method for

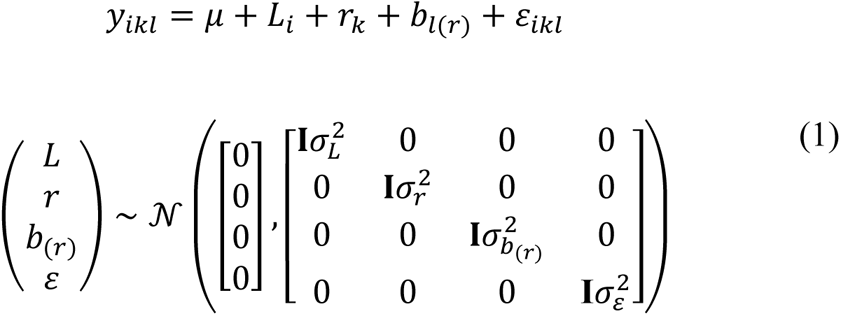

where *y*_*ikl*_ denotes the phenotypic record of the *i^th^* genotype in the *l^th^* block within the *k^th^* replication, *µ* is the overall mean, *L*_*i*_ is the random effect of the *i^th^* genotype, *r*_*k*_ is the random effect of the *k^th^* replication, *b*_*l*(*r*)_is the random nested effect of the *l^th^* block within the *k^th^* replication and *ε*_*ikl*_ is the corresponding error term. These terms are considered as independently and identically distributed outcomes, following a normal distribution with a covariance structure based on the identity matrix **I** and scaled by their corresponding variance component σ^2^.

### 2.4 Allocation strategies

From the initial 220 genotypes, this study focused only on 186 Colombian cultivars. Thus, a total of 1,116 genotype-in-environment combinations (or phenotypes) resulted from the combination of the 186 Colombian genotypes observed at the six different environments. In practical scenarios, budgetary constraints imply that only a subset of these potential phenotypes can be realistically observed in the field. Therefore, strategies are essential in the allocation of genotypes across various environments. The goal is to create a calibration set comprising phenotypic records that maximize the accuracy of prediction models. This is key for obtaining accurate estimates of the performance of genotypes that are not observed in a set of environments. In this study, allocation optimization was executed by classifying the genotypes into one of the following categories:

#### Non-overlapping Genotypes (Genotypes seen only in one environment)

In this category, genotypes are randomly assigned to specific environments in such a way that they are unique and observed only once across environments. For the case of study, this implies that each of the six environments would host 31 unique genotypes (186/6 = 31), guaranteeing that all genotypes are assigned to one environment only with no overlapping across environments.

#### Overlapping Genotypes (Genotypes seen in all the environments)

Some genotypes that exhibit consistent performance across various environments or that are of particular interest for research purposes can be allocated in all six environments. This overlapping ensures that these genotypes are evaluated under diverse conditions, assessing their adaptability and stability across environments.

Let **A** and **B** represent, respectively, the number of non-overlapping and overlapping genotypes, and **T** the number of environments or trials. For a given ‘**A**/**B**’ design, the number of required phenotypes is (**A**+**B**)×**T**, the number of unique genotypes tested is **A**×**T**+**B** and the average number of replicates/phenotypes per genotype is (**A**+**B**×**T**)/(**A**+**B**). Assuming a fixed budget for 186 plots and uniform plot costs in all the environments, two extreme allocation strategies arise.

#### Testing all the genotypes only once (31/0)

In this scenario, 31 unique genotypes are assigned and phenotyped at each environment (**A** = 31). Hence, there is no overlapping of genotypes between environments (**B** = 0). The number of phenotypes is (31+0)×6 = 186, the number of unique tested genotypes is 186 and the number of phenotypes per genotype is (31+0×6)/(31+0) = 1 phenotype/genotype.

#### Testing same genotypes in all environments (0/31)

Here, a subset of 31 unique genotypes from the original set of 186 is phenotyped in all the environments with no specific or non-overlapping genotypes at each environment (**A** = 0) and all the 31 being overlapped genotypes (**B** = 31). The number of phenotypes is still (0+31)×6 = 186 but the number of unique genotypes is reduced to (0 × 6)+31 = 31, and the number of phenotypes per genotype is increased to (0+31×6)/(0+31) = 6 phenotypes/genotype.

In addition, we can construct mixed designs by adding one overlapping genotype and removing one non-overlapping genotype from each environment. For instance, the design 16/15 is composed of 16 non-overlapping genotypes per environment and 15 overlapping genotypes phenotyped in the six environments. In this case, the number of phenotypes remains the same (16+15) × 6 = 186 but the number of unique genotypes is 16×6+15 = 111 and the average phenotypes per genotype is (16+15×6)/(16+15) = 3.42.

Starting from the full non-overlapping design (31/0), nine designs resulted after increasing the number of overlapping or common genotypes (**B**) by four with respect to the previous design, except for the first case where only three genotypes were added to **B**. These designs are 31/0, 28/3, 24/7, 20/11, 16/15, 12/19, 8/23, 4/27, and 0/31. The set of phenotypes or genotype-in-environment combinations observed in fields, (A+B)×T = 186, constitutes the calibration set for training models to predict the phenotypes of the remaining unobserved genotype-in-environment combinations (1,116 - 186 = 930 combinations). A visual representation of the designs is shown in Figure 1.

**Figure 1.**
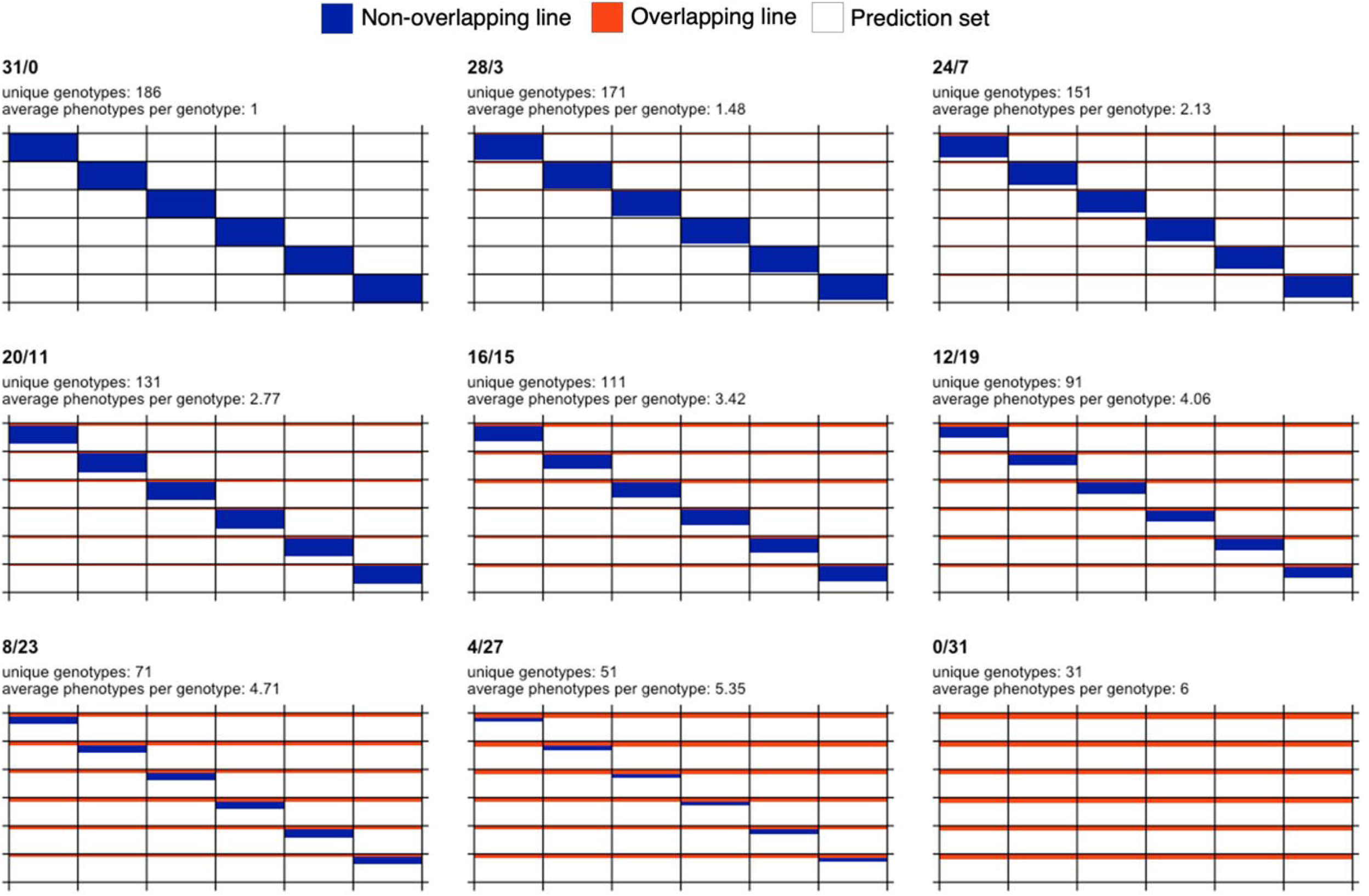
Graphical representation of sparse testing designs for a fixed budget and for a data set comprising 186 phenotypes and six environments. Each subplot represents a different design. The design name, number of unique genotypes and average phenotypes per genotype are shown at the top of each subplot. Rows and columns represent genotypes and environments, respectively. Blue color indicates that a genotype is observed in only one environment, while orange color indicates overlapping genotypes observed in all the six environments. White color is used to represent phenotypes not used in the calibration set and potentially forming the prediction set (155 phenotypes per environment or equivalently 930 phenotypes across environments).

To test the allocation of resources in terms of predictive ability, not only regarding the proportion of non-overlapping and overlapping genotypes, but also with respect to different budget constraints, additional sample sizes were considered for composing training (168, 144, 120, 96, and 72 phenotypes). All calibration set designs are shown in Table 1, where the columns in the middle represent different designs (compositions) and the rows indicate the different calibration sizes. Ten random partitions or replicates were considered to evaluate the predictive ability of the different designs (**combination of set size and composition**). To ensure a fair comparison among sparse designs, the size of the prediction set was fixed to 930 (83.3%) for all these. This entails that some genotype-in-environment combinations were excluded from consideration in rows 2 to 6, neither included in either the calibration or the prediction set.

**Table 1.** Resource allocation designs. The first column indicates the size of the calibration set across environments (relative size with respect to the total potential number of combinations). The last two columns show, the size of the prediction set (constant) and the number (and proportion) of phenotypes discarded from the calibration set. All the designs in the same row share a common calibration set size while the designs in the same column share a common number of non-overlapping genotypes.

| # Calibration | Designs |  |  |  |  |  |  |  |  | # Prediction | # Unused |
| --- | --- | --- | --- | --- | --- | --- | --- | --- | --- | --- | --- |
| 186 (16.7%) | 31/0 | 28/3 | 24/7 | 20/11 | 16/15 | 12/19 | 8/23 | 4/27 | 0/31 | 930 (83.3%) | 0 (0.0%) |
| 168 (15.1%) |  | 28/0 | 24/4 | 20/8 | 16/12 | 12/16 | 8/20 | 4/24 | 0/28 | 930 (83.3%) | 18 (1.6%) |
| 144 (12.9%) |  |  | 24/0 | 20/4 | 16/8 | 12/12 | 8/16 | 4/20 | 0/24 | 930 (83.3%) | 42 (3.8%) |
| 120 (10.8%) |  |  |  | 20/0 | 16/4 | 12/8 | 8/12 | 4/16 | 0/20 | 930 (83.3%) | 66 (5.9%) |
| 96 (8.6%) |  |  |  |  | 16/0 | 12/4 | 8/8 | 4/12 | 0/16 | 930 (83.3%) | 90 (8.1%) |
| 72 (6.5%) |  |  |  |  |  | 12/0 | 8/4 | 4/8 | 0/12 | 930 (83.3%) | 114 (10.2%) |

### 2.5 Predictive models

Three different GP models, based on random effects and modeled via covariance structures, were calibrated and used to predict the testing sets. The first model **M1** is considered the base model, and it assumes independent and identically distributed (IID) outcomes among its components. The second model, **M2**, leverages molecular marker information to determine relationships between pairs of genotypes. While **M1** and **M2** are main effects models, the third model **M3** introduces the genotype-by-environment (G×E) interaction term. The model’s performance was assessed using the Pearson’s correlation between the observed and the predicted values within each environment. The traits sucrose accumulation (SA) and tons of cane per hectare (TCH) were analyzed separately.

#### M1: Environment and genotype main effects

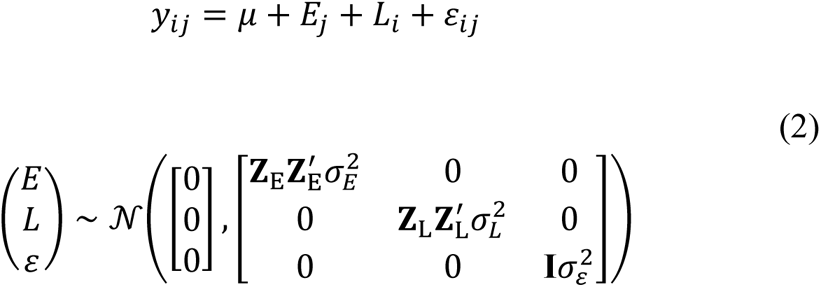

where *y*_*ij*_ is the adjusted phenotypic observation of the target trait in the *j^th^* environment (*j=*1,2,…,6) for the *i^th^* genotype (*i=*1,2,…,186)*, µ* is the overall mean, *E*_*j*_ is the random effect of the *j^th^* environment, *L*_*i*_ is the random effect of the *i^th^* genotype, and *ε*_*ij*_ is the corresponding error term. *E*_*j*_, *L*_*i*_, and *ε*_*ij*_ are assumed to be independent and identically distributed outcomes from a normal density. **Z**_E_ (*n* × 6) and **Z**_L_ (*n* × 186) are the design matrices connecting phenotypes with environments and genotypes, respectively, and **I** (*n* × *n*) is the identity matrix, with *n* being the number of total observations (*n* = 186 × 6). The variance component for each term is denoted by σ^2^. In this model, the genotypes are assumed to be independent, and therefore, no information can be borrowed from phenotyped individuals to their nonphenotyped relatives.

#### M2: Environment, genotype, and genomic markers main effects

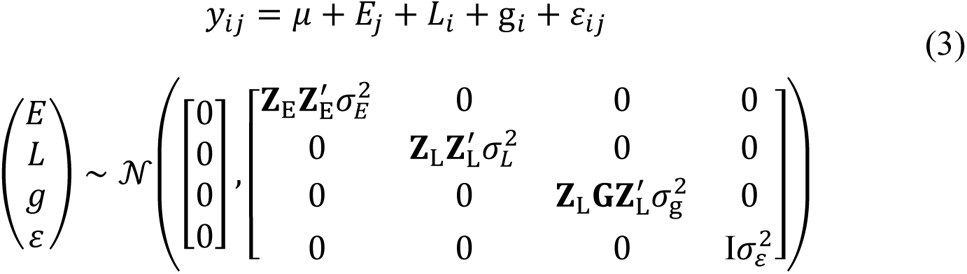

where all the terms in common with **M1** have the same meaning, g_*i*_ is the genomic effect based on maker SNPs for the *i^th^*genotype, and **G** (186 × 186) is the kinship matrix describing genomic similarities between pairs of individuals (Van Raden, 2008). Both *L*_*i*_ and g_*i*_ terms capture information about the genotype but in the latter, genomic information is used to relate phenotyped and non-phenotyped individuals. The genotype effect *L*_*i*_ is not removed from the model to account for model miss-specification and imperfect information (genetic variability that cannot be explained through SNP markers only).

#### M3: Environment, genotype, and genomic markers main effects, and Genotype × Environment interaction

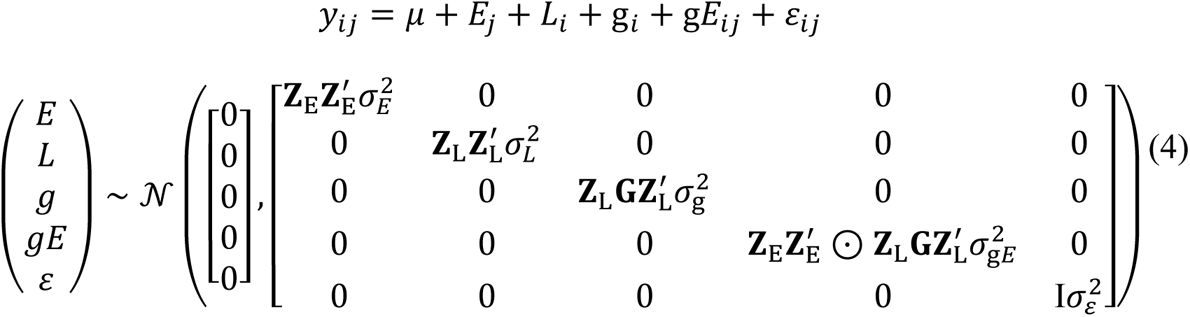

where all the terms in common with **M2** remain the same, and the added term g*E*_*ij*_ represents the effect of the interaction between the *j^th^* environment and *i^th^* genotype (Jarquin *et al.,* 2014), ⊙ is the element-wise multiplication or Hadamard product operation between two matrices.

### 2.6 Cross Validation and performance metrics

In the context of plant breeding, particularly in the GP domain, the evaluation of models often involves the use of cross-validation schemes (Jarquin et al., 2021). One widely used method is the K-fold cross-validation, which helps to assess the performance of predictive models. K-fold cross-validation involves splitting the dataset into K subsets or folds. The process then iterates K times, each time using one of the K subsets as the prediction set while the remaining K-1 subsets are used for calibrating the predictive models. This approach provides a robust way to assess the model’s performance. It returns predicted values for all the datapoints in a fold without any of them being present in the calibration set.

The process of partitioning the dataset into folds is not trivial and might include some constraints. In this study, we included features of two common cross-validation schemes: CV2 and CV1. In the basic cross-validation scheme, known as CV2, the dataset is randomly partitioned without any specific constraints. This means that the model may predict the phenotype of genotypes that have been observed in other environments. While this approach is informative, it doesn’t guarantee that the model will predict unseen genotypes, as some genotypes may appear in both the calibration and the prediction sets due to random partitioning. In CV1, a specific constraint is applied: all phenotypes of the same genotype must be placed in the same fold. This ensures that when evaluating the model, none of the observations of a genotype in any environment are present in the calibration set. This constraint guarantees that the models will predict an unseen genotype, as all observations of that genotype are in the prediction test.

The allocation designs constitute the spectrum between CV2 and CV1, with 31/0 being the pure CV2 and 0/31 being the CV1 scheme. The higher the number of overlapping genotypes, the closer to the CV1 scheme. All the designs except those with 0 non-overlapping genotypes are CV2-like schemes. In the context of predictive modeling, the CV1 scheme poses a greater challenge, especially for models that do not incorporate genomic information and treat genotypes as independent entities. This challenge arises because the calibration set does not provide any useful insights during the model training process for predicting a specific, new, and unseen genotype.

To evaluate the performance of predictive models in this study, Pearson’s correlation was used as a measure of the association between the adjusted phenotypes and the predicted values. These correlations were computed separately for each environment or trial, reflecting how well the models performed when predicting the genotypes in specific environmental conditions. Averaged correlations from the 10 replications or partitions were obtained.

To further interpret the results of models’ performance, we computed the percentage of variance explained by each term in the different fitted models after conducting a full dataset analysis (i.e., with no missing values).

### 2.7 Software

Genomic prediction analyses were computed in the R environment (R Development Core Team, 2023) and the models were fitted using the BGLR package (Pérez and de los Campos, 2014). All models were fitted using 12,000 Markov Chain Monte Carlo iterations, incorporating a burn-in phase for the initial 2,000 iterations and a thinning factor of 5.

## 3 Results

### 3.1 Variance components

The percentage of variance explained by each term for SA and TCH is shown in Table 2. For both traits, the environmental component captured the largest proportion of variability. However, there are significant differences between traits, with 46%-48% of the total variance explained for SA while 80%-82% for TCH. Examining the decomposition of genetic variance, we found that for SA the variance explained by genotype in model 1 (56.2% of the within-environments variance) is higher than the sum of the variance explained by genotype and genomic markers (12.9% + 34.5% = 47.4%) in model 2. A similar pattern was observed for TCH (34.9% in model 1 and 23.0% + 12.1% = 35.1% in model 2). Conversely, the variance explained by the interaction between genomic markers and environment is orthogonal to the other components, explaining the variability that cannot be captured by the main effects, thus reducing the residual variance.

**Table 2.** Across and within environments percentage of explained variability by each model term for each trait. The first set of columns on the left side depict the percentage for all model terms: main effect of environments (vE), main effect of Genotype (vL), main effect of genomic markers (vg) and the interaction between genomic markers and environments (vgE), as well as the residual or unexplained variability (vR). The second set of columns on the right side depict the within-environment variance, i.e., the relative contribution of each term without considering the variance explained by the main effect of the environments.

| Trait | Model | Across-environments variance (%) |  |  |  |  | Within-environments variance (%) |  |  |  |
| --- | --- | --- | --- | --- | --- | --- | --- | --- | --- | --- |
|  |  | vE | vL | vg | vgE | vR | vL | vg | vgE | vR |
| SA | L+E | 46.7 | 29.9 |  |  | 23.3 | 56.2 |  |  | 43.8 |
|  | L+E+g | 48.1 | 6.7 | 17.9 |  | 27.3 | 12.9 | 34.5 |  | 52.6 |
|  | L+E+g+gE | 47.7 | 6.2 | 18.0 | 15.4 | 12.7 | 11.9 | 34.4 | 29.5 | 24.2 |
| TCH | L+E | 82.2 | 6.2 |  |  | 11.6 | 34.9 |  |  | 65.1 |
|  | L+E+g | 81.3 | 4.3 | 2.3 |  | 12.1 | 23.0 | 12.1 |  | 64.9 |
|  | L+E+g+gE | 80.7 | 4.0 | 2.1 | 4.5 | 8.7 | 20.9 | 10.8 | 23.5 | 44.8 |

### 3.2 Phenotypic correlation between environments

The phenotypic correlation values between environments for both SA and TCH traits are shown in Figure 2. A wide range of positive correlations was found between environments, with no high correlation values between environments sharing locations except in the case of SA in Balsora environments (r = 0.822). Correlation values for Balsora environments with other locations were higher than those between Taula and Porvenir for SA. For TCH, we found lower correlations, with the maximum values observed between Taula 2019 and Porvenir environments (r = 0.621 and r = 0.588 for 2020 and 2021, respectively).

**Figure 2.**
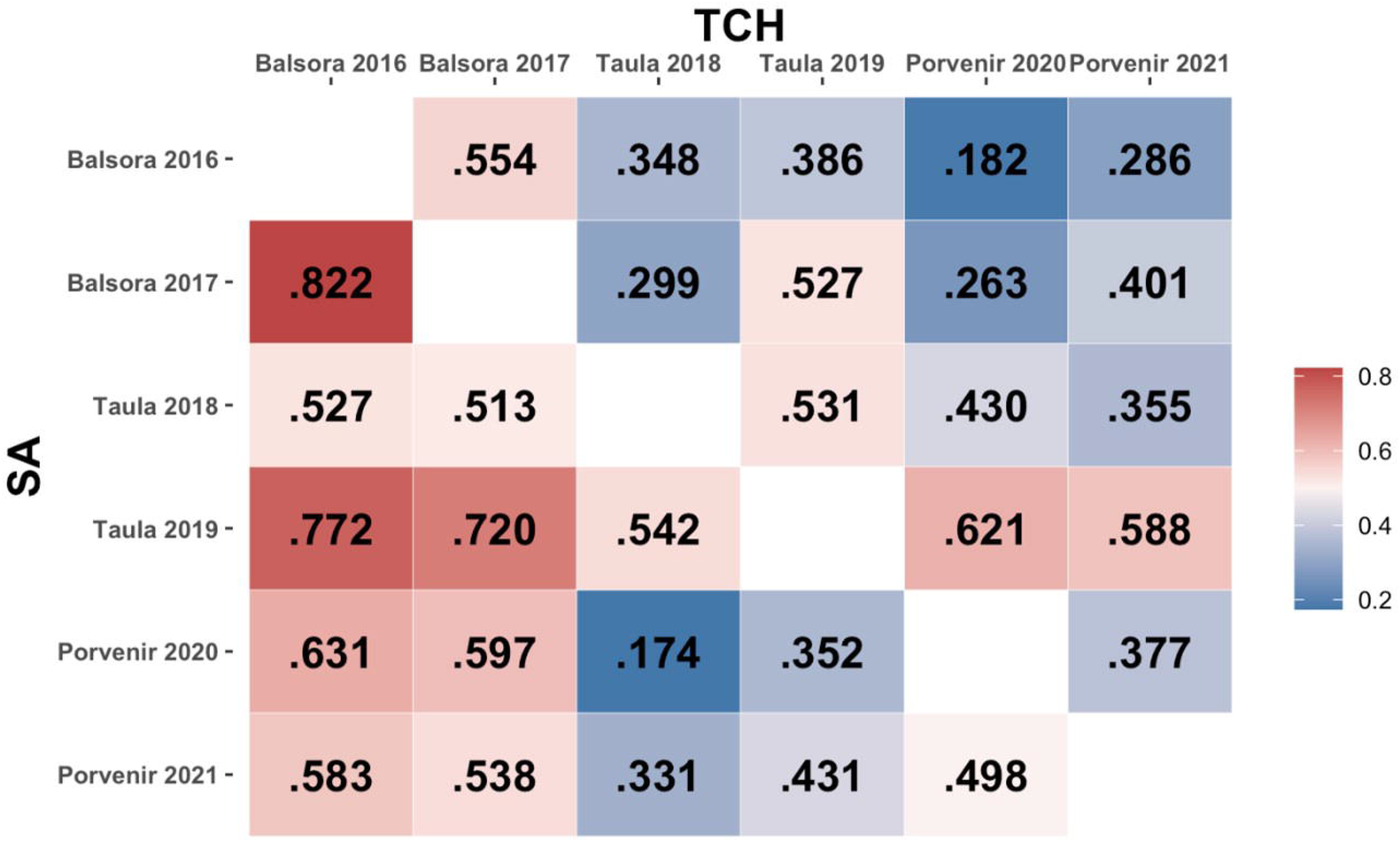
Phenotypic correlation between environments. The lower and upper triangular matrices represent, respectively, SA and TCH traits. Color reference is missing.

Figure 3 presents the accuracy achieved at each environment, as well as the overall mean accuracy. The original values used to compute the means, as well as the values for scenarios with reduced calibration set are presented in Supplementary Table 1. The accuracy was generally higher when predicting SA compared to TCH for models M2 and M3 (M1 returned low results due to the lack of information to connect calibration and testing sets). As a general trend, the accuracy decreases as the number of overlapping genotypes increases. However, within the SA trait, the maximum accuracy was attained at the ‘28/3’ allocation design. Furthermore, a local optimum was identified at the ‘12/19’ allocation design, indicating that specific combinations of overlapping and non-overlapping genotypes can lead to improved predictive performance.

**Figure 3.**
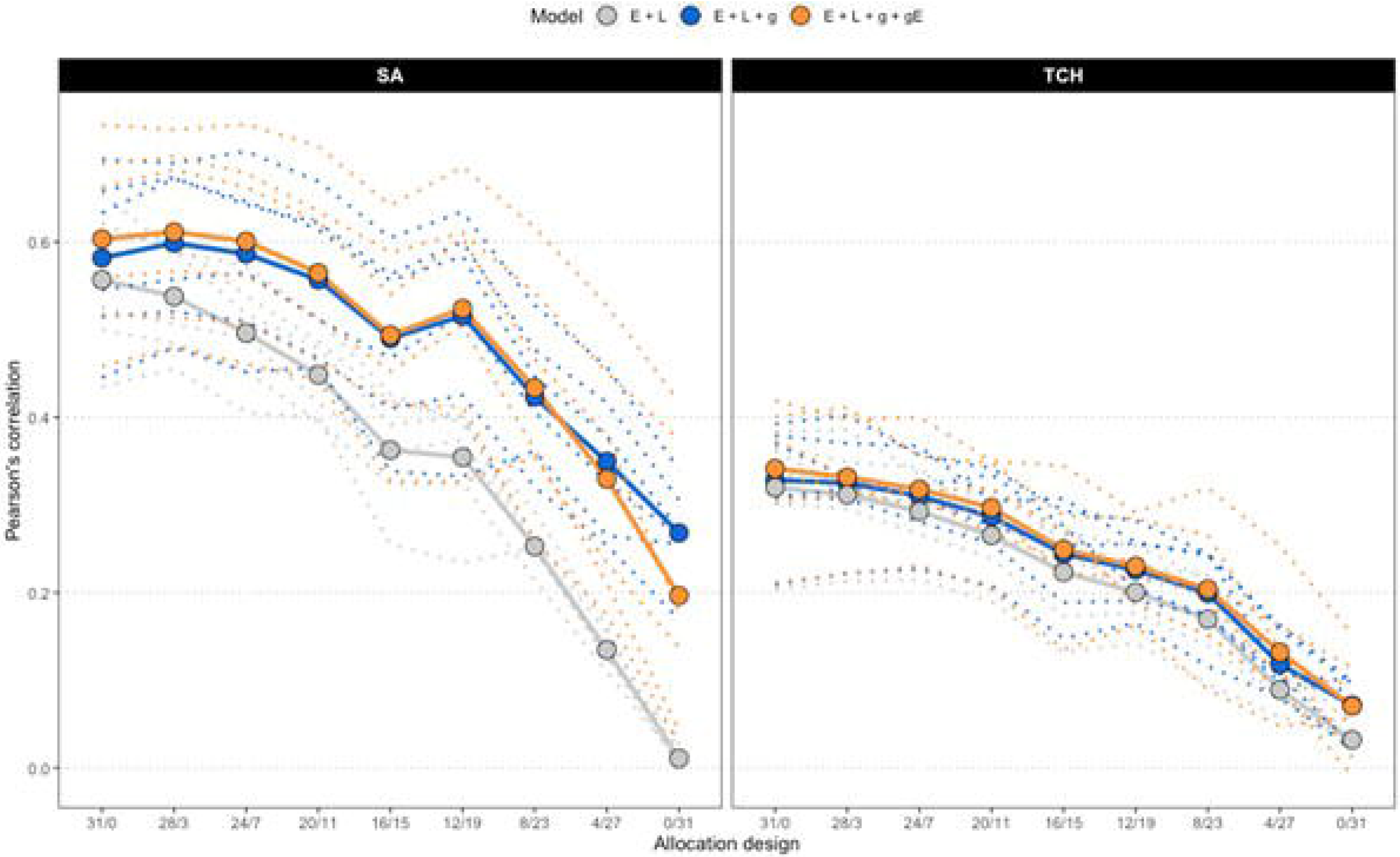
Average accuracy of models. Due to the increasing number of combinations, only the results for the allocation designs with the largest calibration set size (186) are shown for SA (left) and TCH (right). The *x*-axis represents the allocation designs, and the *y*-axis represents the accuracy measured as the Pearson’s correlation between predicted and observed values. Color denotes models. Dotted lines represent one specific environment. Solid lines and highlighted points represent the average accuracy across six environments.

Regarding the models used in the study, it was found that M3, which accounts for genotype-by-environment (G×E) interactions, generally outperformed the other models, but there were not significant differences with respect to M2, as stated in Supplementary Table 2. Additionally, as expected, the base model M1 consistently showed the lowest predictive accuracy across all scenarios, with the performance gap widening as the number of overlapping genotypes increased. Notably, the study consistently achieved the best predictions in specific environments, namely E1 (Balsora 2016), E2 (Balsora 2017), and E4 (Taula 2019), as can be noted in Supplementary Table 1. Figure 4 shows the average accuracy obtained across all calibration sizes and designs. In alignment with the results from Figure 3, we found that accuracy tends to decline as the number of non-overlapping genotypes decreases. This trend holds true for all models and designs for TCH, with a few exceptions for SA, specifically ‘12/19’, ‘8/20’, and ‘8/12’ for models M2 and M3.

**Figure 4.**
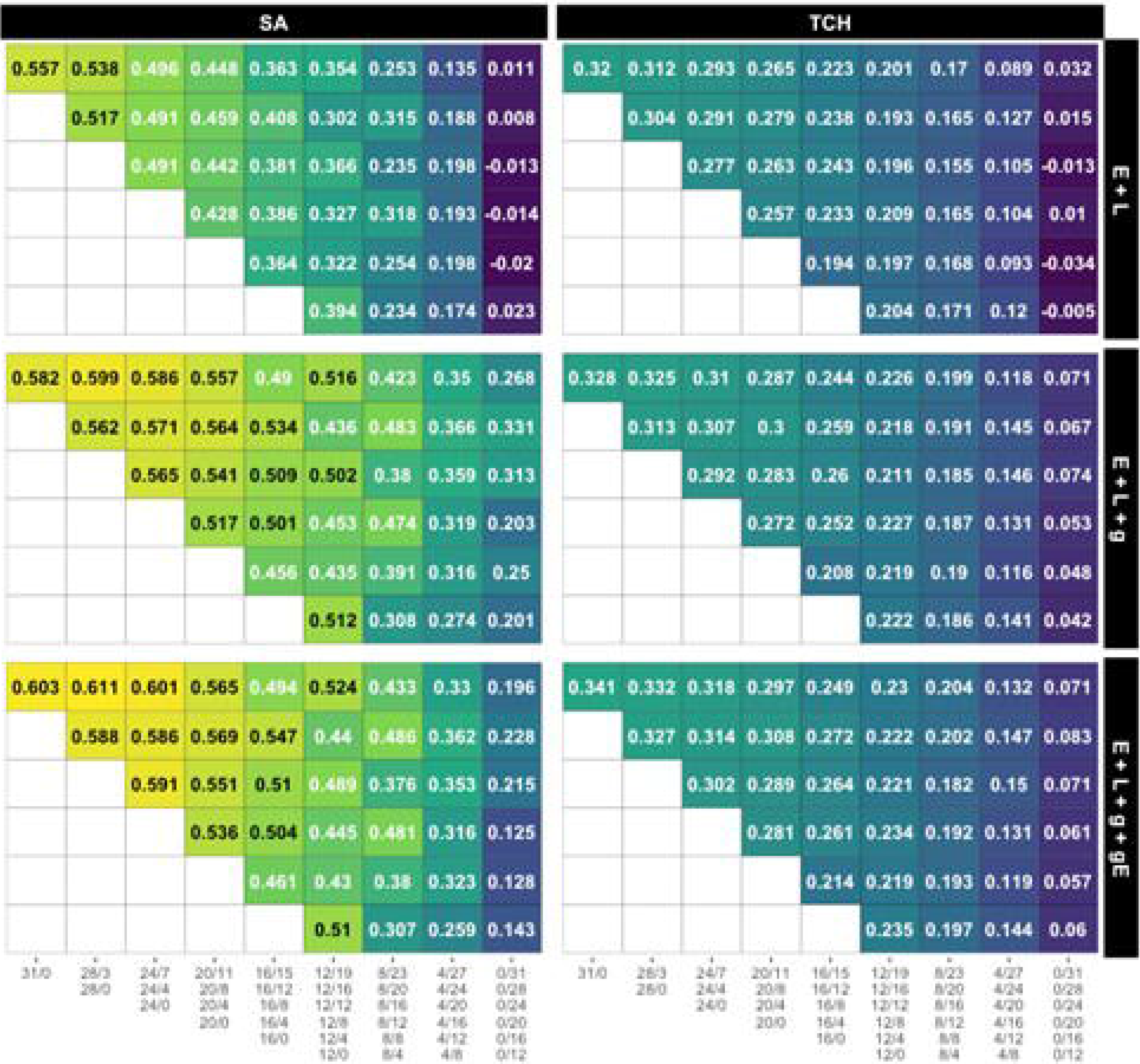
Average accuracy for all designs and calibration sizes. Each subplot represents the accuracy obtained by each combination of model and trait. Each grid within a subplot represents the accuracy for a specific design and calibration size, with yellow, green and blue colors representing high, medium and low accuracy values, respectively. Grids in the same row share the calibration size while grids in the same column share a common number of non-overlapping genotypes. The specific designs are presented below the columns.

In contrast to the impact of decreasing the number of non-overlapping genotypes, reducing the calibration size (i.e., the number of plots) by allocating a lower number of overlapping genotypes appears to have a minimal effect on accuracy, as shown in the main diagonals (*i* = *j*) of Figure 4. Remarkably, in models M2 and M3, the accuracy for the ‘12/0’ design surpasses that of all other designs within the ‘12/B’ family, except for the one with the larger calibration size and, therefore, number of overlapping genotypes (’12/19’). This suggests that similar accuracy levels could be achieved by either decreasing the calibration size and using fewer overlapping genotypes or by maintaining the calibration size while evaluating more non-overlapping genotypes.

## 4 Discussion

Modern plant breeding strives to find a balance between increasing genetic gain and reducing resource allocation. Given fixed financial resources, it is often challenging to determine how to allocate a predetermined number of genotypes for evaluation in targeted environments. To address this, optimization of resources in multi-environment trials has become a routine practice. Among the strategies employed, the sparse testing allocation scheme with genomic prediction has been promising. This approach could identify the minimal number of candidate genotypes required for evaluation in METs and strategize their distribution in replicated or unreplicated field designs to achieve maximum genetic gains. In this study, we explored different sparse testing designs to determine the optimal calibration size and the trade-off threshold between prediction accuracy and selection intensity in sugarcane breeding.

One of the most notable trends is that CV1-like schemes (‘0/B’) pose significant challenges for M1: E + L. This phenomenon could be attributed to the fact that the predicted value of a genotype in each environment depends on the information of that exact genotype in other environments, which is not available in CV1 scenarios. By borrowing information from related genotypes through genomic information, M2 and M3 can extract patterns from the phenotypes of related genotypes and therefore, have a reasonable performance on CV1 scenarios. Similar results have been reported in a previous study by Atanda et al. (2022), where genomic and pedigree relationship between individuals can track segregating Quantitative Trait Loci, thus, explaining a large proportion of the genetic variance in a population.

On the other extreme, for the CV2-like schemes (‘A/0’) there were no notable differences in the predictive abilities of the models, even though they showed high predictive power (Figure 4). This is because the environmental effects are confounded with the genotype effect such that a single observation of a genotype in an environment is enough to predict its performance in other environments. Therefore, predictive ability strongly depends on the phenotypic information of tested individuals. Incorporation of genomic information explained a low genomic variance (SA: 17.9%, and TCH: 2.3%) in M2 (Table 2) and the model accounting for genotype by environment interaction in M3 captured a small information from related genotypes evaluated in correlated environments (SA: 15.4%, and TCH: 4.5%). This result implies a high correlation between the environments, given the observed small genotype differentiation across them. As such, there is a need to redefine the target population of environments to avoid resource wastage during multi-environment trials. These environments can be classified using important environmental factors and consolidated into mega-environments for efficient resource allocation (Resende et al., 2021).

The best accuracy is consistently obtained in scenarios with high ‘A’ value. The larger the number of different genotypes observed at least once, the higher the performance achieved. This suggests that G×E interactions are small and, therefore, phenotyping a genotype in one environment provides useful insights for the prediction of the remaining environments. However, this might seem contradictory to the findings in Table 2, where a significant portion of the Within-environment variance was attributed to the G×E effect. It is crucial to interpret those results with caution because, while all the 1,116 available phenotypes where used to calibrate the model that estimate the variances, predictive models were trained using a maximum of 186 phenotypes.

Moreover, the slight gain in accuracy obtained by M3, which accounts for G×E interactions occurs in the scenarios with more non-overlapping genotypes. These findings indicate a potential cost saving opportunity can be achieved by allocating the genotypes to non-overlapping environments, that is, observing each genotype once across environment. This could decrease the cost of METs significantly in sugarcane breeding, as overlapping genotypes in some environments showed no gains in genomic prediction. This result is similar to the findings of Jarquin et al. (2020) in maize where marginal gains were observed even when overlapping genotypes were increased significantly. The plausible explanation is either the environments share strong environmental similarly or the traits exhibit low response to genotype by environment interaction. In that sense, Balsora, Taula and Porvenir locations are characterized by having the same soil taxonomy with soils from vertisol, inceptisol and mollisol orders, which have a fine, dry, and soils from deep to moderately deep (Carbonell et al., 2001), making them locations with similar behavior. However, even though the locations (E1 to E6) have similar soil profiles, Taula (E3 and E4) and Porvenir (E5 and E6) also share a lower drainage capacity when compared with Balsora (E1 and E2), affecting the performance of the planted varieties, impacting positively the correlations of sucrose and TCH among locations.

The breeding program at Cenicaña involves making decisions after 12 years of selection based on the phenotypic information collected during the early stages of selection (three stages). The multi-environmental trials (METs) provide information of genotype adaptability to the target population of environments of Colombia (CENICAÑA, 1995). It is becoming important to integrate the allocation of phenotypes in plant breeding programs to optimize limited resources and increase genetic gain.

Given the availability of genomic information during the breeding process, this can be leveraged to either evaluate more genotypes by increasing selection intensity or maximize the genetic gains with a fixed plot unit cost in sparse testing. As such, findings from this study in the case of sugar cane indicated that a cost saving opportunity could be achieved by overlapping a small number of genotypes in all environments while allocating the remaining genotypes to different environments. In addition, the use of a genomic prediction model that incorporates G×E interaction (M3) allowed to capture genetic information among related genotypes in other environments to improve predictive ability.

The assumption of independence between genotypes held in M1 leads to biased models that perform poorly in comparison with the models that integrate genomic information to connect these. However, there is a clear trend to decrease accuracy with decreasing calibration set sizes. While the best accuracy is obtained with the full calibration set, there are designs with reduced calibration sizes that equal or even overpass the equivalent designs with larger calibration sizes. This is found especially in SA trait. Therefore, we can obtain similar accuracy values by reducing the budget or keeping the same budget but increasing the number of unique genotypes evaluated by increasing the number of non-overlapping genotypes. This is similar to what has been obtained in maize, wheat, and soybean sparse testing allocation designs (Jarquin et al., 2020; Crespo-Herrera et al., 2021; Persa et al., 2023) but the difference we observed in sugarcane is that increasing the number of overlapping genotypes did not resulted in an increased predictive ability. This might also be due to the reduced training set sizes of our dataset compared with these studies. Interestingly, there are major differences in the average accuracy between traits. In the optimum scenarios, ‘31/0’ and ‘28/3’, the accuracy values for SA are 0.603 and 0.611, while for TCH. are 0.341 and 0.332. SA is a more heritable trait with an extended genetic control while TCH trait is highly dependent on environmental conditions, specifically precipitation. Moreover, these differences might be caused by the phenotyping procedures.

## 5 Conclusions

In this study, we investigated the potential of genomic prediction for sparse testing resource allocation in sugarcane multi-environmental trials. It involved the utilization of different ratios of NOG/OG sugarcane clones and genome-based models, including G×E, to capture more genetic variability other than the main genomic effects. The results obtained indicated that genomic prediction models that incorporated G×E had the highest predictive response compared to other models in all allocation design scenarios. However, the advantage of the genomic model with G×E decreased with increasing NOG where highest predictive values were obtained. While this study focused on maximizing the genetic gain with a fixed cost per phenotype in sparse testing, the results showed that reducing the sample size of the genotypes assigned to environments (NOG) decreased the accuracy of genomic prediction. The trend decreased further with increasing overlapping genotypes evaluated across environments. This indicates that very few overlapping genotypes are needed across environments. This was attributed to a high environmental effect on the traits and moderate phenotypic correlation between environments. Generally, the results from this study showed that models including G×E can reduce resource allocation for phenotyping by up to 83% or increase the testing capacity by fivefold for multi-environmental trials in sugarcane. Therefore, sparse testing with genomic prediction is a promising strategy for maximizing genetic with fixed phenotyping cost in a breeding program. Premised on large environmental variance for the traits, we recommend the use of environmental factors to define TPEs to avoid investing limited resources in correlated environments without corresponding increase in genetic gain.

## 7 Author Contributions

JGA: Writing – original draft, Conceptualization, Methodology, Formal analysis, Visualization; PA: Writing – review & editing; FSA: Writing - review & editing, Conceptualization, Resources, Data curation; JHTM: Writing – review & editing, Resources; JJR: Writing – review & editing, Conceptualization, Resources, Data curation; RPS: Writing - review & editing; JIS: Writing - review & editing, Resources; DJ: Conceptualization, Methodology, Formal analysis, Supervision, Project administration,.

## 9 Acknowledgments

The authors would like to thank the mills and growers who have supported Cenicaña for more than 46 years. Additionally, we would like to give special thanks to the plant breeders at Cenicaña who have been actively releasing commercial varieties in Colombia.

## 10 Data Availability Statement

The datasets and pipeline generated for this study can be found in the following link: https://uflorida-my.sharepoint.com/:f:/g/personal/jhernandezjarqui_ufl_edu/Ele_tW5RgC5PrfRHWnet5xsBzgOHCLzCZ_Yxz8SftLUv-Q?e=WhhHaq

## Supplementary Material

**Supplementary Table 1.** Average Pearson’s correlation values between observed and predicted phenotypic values for 10 replicates for different sparse testing designs and environments (E1: Balsora 16, E2: Balsora 17, E3: Taula 18, E4: Taula 19, E5: Porvenir 20, E6: Porvenir 21).

| Model | Trait | SS | Design | E1 | E2 | E3 | E4 | E5 | E6 | Mean |
| --- | --- | --- | --- | --- | --- | --- | --- | --- | --- | --- |
| M1: E + L | SA | 31 | 31/0 | 0.65 | 0.62 | 0.52 | 0.5 | 0.43 | 0.61 | 0.56 |
| M1: E + L | SA | 31 | 28/3 | 0.59 | 0.59 | 0.51 | 0.48 | 0.46 | 0.6 | 0.54 |
| M1: E + L | SA | 31 | 24/7 | 0.57 | 0.52 | 0.49 | 0.45 | 0.41 | 0.54 | 0.5 |
| M1: E + L | SA | 31 | 20/11 | 0.51 | 0.48 | 0.42 | 0.4 | 0.4 | 0.49 | 0.45 |
| M1: E + L | SA | 31 | 16/15 | 0.43 | 0.39 | 0.36 | 0.32 | 0.25 | 0.42 | 0.36 |
| M1: E + L | SA | 31 | 12/19 | 0.39 | 0.4 | 0.37 | 0.32 | 0.23 | 0.4 | 0.35 |
| M1: E + L | SA | 31 | 8/23 | 0.29 | 0.26 | 0.25 | 0.21 | 0.25 | 0.25 | 0.25 |
| M1: E + L | SA | 31 | 4/27 | 0.16 | 0.15 | 0.13 | 0.11 | 0.12 | 0.13 | 0.13 |
| M1: E + L | SA | 31 | 0/31 | 0 | 0 | 0.02 | 0.01 | 0.03 | 0 | 0.01 |
| M1: E + L | TCH | 31 | 31/0 | 0.36 | 0.37 | 0.2 | 0.3 | 0.3 | 0.38 | 0.32 |
| M1: E + L | TCH | 31 | 28/3 | 0.32 | 0.35 | 0.21 | 0.31 | 0.29 | 0.39 | 0.31 |
| M1: E + L | TCH | 31 | 24/7 | 0.29 | 0.34 | 0.21 | 0.3 | 0.27 | 0.33 | 0.29 |
| M1: E + L | TCH | 31 | 20/11 | 0.26 | 0.27 | 0.19 | 0.32 | 0.24 | 0.32 | 0.27 |
| M1: E + L | TCH | 31 | 16/15 | 0.23 | 0.28 | 0.13 | 0.26 | 0.17 | 0.27 | 0.22 |
| M1: E + L | TCH | 31 | 12/19 | 0.23 | 0.22 | 0.14 | 0.18 | 0.18 | 0.25 | 0.2 |
| M1: E + L | TCH | 31 | 8/23 | 0.22 | 0.2 | 0.1 | 0.15 | 0.14 | 0.21 | 0.17 |
| M1: E + L | TCH | 31 | 4/27 | 0.08 | 0.11 | 0.06 | 0.08 | 0.08 | 0.13 | 0.09 |
| M1: E + L | TCH | 31 | 0/31 | 0.02 | 0.03 | 0.04 | 0.03 | 0.03 | 0.05 | 0.03 |
| M2: E + L + g | SA | 31 | 31/0 | 0.69 | 0.66 | 0.55 | 0.51 | 0.45 | 0.63 | 0.58 |
| M2: E + L + g | SA | 31 | 28/3 | 0.69 | 0.67 | 0.56 | 0.52 | 0.48 | 0.67 | 0.6 |
| M2: E + L + g | SA | 31 | 24/7 | 0.7 | 0.64 | 0.56 | 0.51 | 0.45 | 0.65 | 0.59 |
| M2: E + L + g | SA | 31 | 20/11 | 0.67 | 0.62 | 0.51 | 0.47 | 0.46 | 0.61 | 0.56 |
| M2: E + L + g | SA | 31 | 16/15 | 0.6 | 0.56 | 0.47 | 0.41 | 0.34 | 0.56 | 0.49 |
| M2: E + L + g | SA | 31 | 12/19 | 0.63 | 0.6 | 0.52 | 0.43 | 0.33 | 0.58 | 0.52 |
| M2: E + L + g | SA | 31 | 8/23 | 0.53 | 0.48 | 0.41 | 0.32 | 0.36 | 0.45 | 0.42 |
| M2: E + L + g | SA | 31 | 4/27 | 0.46 | 0.42 | 0.33 | 0.25 | 0.26 | 0.38 | 0.35 |
| M2: E + L + g | SA | 31 | 0/31 | 0.34 | 0.31 | 0.26 | 0.17 | 0.26 | 0.27 | 0.27 |
| M2: E + L + g | TCH | 31 | 31/0 | 0.37 | 0.38 | 0.21 | 0.31 | 0.31 | 0.39 | 0.33 |
| M2: E + L + g | TCH | 31 | 28/3 | 0.33 | 0.37 | 0.22 | 0.32 | 0.31 | 0.4 | 0.33 |
| M2: E + L + g | TCH | 31 | 24/7 | 0.31 | 0.37 | 0.23 | 0.32 | 0.28 | 0.36 | 0.31 |
| M2: E + L + g | TCH | 31 | 20/11 | 0.28 | 0.3 | 0.21 | 0.33 | 0.26 | 0.34 | 0.29 |
| M2: E + L + g | TCH | 31 | 16/15 | 0.25 | 0.31 | 0.15 | 0.28 | 0.19 | 0.29 | 0.24 |
| M2: E + L + g | TCH | 31 | 12/19 | 0.26 | 0.26 | 0.17 | 0.2 | 0.19 | 0.28 | 0.23 |
| M2: E + L + g | TCH | 31 | 8/23 | 0.25 | 0.24 | 0.12 | 0.17 | 0.17 | 0.24 | 0.2 |
| M2: E + L + g | TCH | 31 | 4/27 | 0.11 | 0.16 | 0.08 | 0.1 | 0.1 | 0.16 | 0.12 |
| M2: E + L + g | TCH | 31 | 0/31 | 0.1 | 0.1 | 0.02 | 0.03 | 0.09 | 0.1 | 0.07 |
| M3: E + L + g + gE | SA | 31 | 31/0 | 0.73 | 0.69 | 0.56 | 0.52 | 0.46 | 0.66 | 0.6 |
| M3: E + L + g + gE | SA | 31 | 28/3 | 0.73 | 0.7 | 0.57 | 0.51 | 0.48 | 0.68 | 0.61 |
| M3: E + L + g + gE | SA | 31 | 24/7 | 0.73 | 0.68 | 0.56 | 0.51 | 0.46 | 0.66 | 0.6 |
| M3: E + L + g + gE | SA | 31 | 20/11 | 0.71 | 0.64 | 0.51 | 0.46 | 0.44 | 0.62 | 0.56 |
| M3: E + L + g + gE | SA | 31 | 16/15 | 0.64 | 0.59 | 0.45 | 0.41 | 0.33 | 0.54 | 0.49 |
| M3: E + L + g + gE | SA | 31 | 12/19 | 0.69 | 0.61 | 0.5 | 0.41 | 0.33 | 0.6 | 0.52 |
| M3: E + L + g + gE | SA | 31 | 8/23 | 0.62 | 0.54 | 0.36 | 0.26 | 0.35 | 0.47 | 0.43 |
| M3: E + L + g + gE | SA | 31 | 4/27 | 0.53 | 0.45 | 0.21 | 0.18 | 0.23 | 0.38 | 0.33 |
| M3: E + L + g + gE | SA | 31 | 0/31 | 0.42 | 0.37 | 0.04 | 0.03 | 0.14 | 0.18 | 0.2 |
| M3: E + L + g + gE | TCH | 31 | 31/0 | 0.37 | 0.42 | 0.21 | 0.31 | 0.34 | 0.4 | 0.34 |
| M3: E + L + g + gE | TCH | 31 | 28/3 | 0.33 | 0.4 | 0.22 | 0.31 | 0.32 | 0.41 | 0.33 |
| M3: E + L + g + gE | TCH | 31 | 24/7 | 0.31 | 0.4 | 0.23 | 0.32 | 0.3 | 0.36 | 0.32 |
| M3: E + L + g + gE | TCH | 31 | 20/11 | 0.27 | 0.35 | 0.2 | 0.32 | 0.29 | 0.35 | 0.3 |
| M3: E + L + g + gE | TCH | 31 | 16/15 | 0.24 | 0.34 | 0.14 | 0.27 | 0.22 | 0.29 | 0.25 |
| M3: E + L + g + gE | TCH | 31 | 12/19 | 0.24 | 0.29 | 0.16 | 0.19 | 0.22 | 0.28 | 0.23 |
| M3: E + L + g + gE | TCH | 31 | 8/23 | 0.19 | 0.32 | 0.09 | 0.15 | 0.21 | 0.27 | 0.2 |
| M3: E + L + g + gE | TCH | 31 | 4/27 | 0.1 | 0.25 | 0.05 | 0.09 | 0.16 | 0.13 | 0.13 |
| M3: E + L + g + gE | TCH | 31 | 0/31 | 0.1 | 0.15 | 0.06 | -0.01 | 0.11 | 0.01 | 0.07 |
| M1: E + L | SA | 28 | 28/0 | 0.62 | 0.55 | 0.51 | 0.42 | 0.41 | 0.59 | 0.52 |
| M1: E + L | SA | 28 | 24/4 | 0.54 | 0.53 | 0.45 | 0.45 | 0.41 | 0.55 | 0.49 |
| M1: E + L | SA | 28 | 20/8 | 0.54 | 0.48 | 0.44 | 0.43 | 0.38 | 0.49 | 0.46 |
| M1: E + L | SA | 28 | 16/12 | 0.47 | 0.44 | 0.36 | 0.37 | 0.36 | 0.44 | 0.41 |
| M1: E + L | SA | 28 | 12/16 | 0.36 | 0.32 | 0.29 | 0.28 | 0.22 | 0.34 | 0.3 |
| M1: E + L | SA | 28 | 8/20 | 0.37 | 0.34 | 0.33 | 0.3 | 0.2 | 0.35 | 0.32 |
| M1: E + L | SA | 28 | 4/24 | 0.21 | 0.18 | 0.19 | 0.16 | 0.2 | 0.18 | 0.19 |
| M1: E + L | SA | 28 | 0/28 | -0.02 | -0.02 | 0.03 | 0.04 | 0.02 | -0.01 | 0.01 |
| M1: E + L | TCH | 28 | 28/0 | 0.33 | 0.34 | 0.2 | 0.32 | 0.28 | 0.36 | 0.3 |
| M1: E + L | TCH | 28 | 24/4 | 0.28 | 0.33 | 0.2 | 0.3 | 0.28 | 0.35 | 0.29 |
| M1: E + L | TCH | 28 | 20/8 | 0.28 | 0.32 | 0.21 | 0.29 | 0.25 | 0.32 | 0.28 |
| M1: E + L | TCH | 28 | 16/12 | 0.24 | 0.25 | 0.16 | 0.27 | 0.22 | 0.28 | 0.24 |
| M1: E + L | TCH | 28 | 12/16 | 0.22 | 0.23 | 0.11 | 0.23 | 0.15 | 0.22 | 0.19 |
| M1: E + L | TCH | 28 | 8/20 | 0.19 | 0.2 | 0.1 | 0.16 | 0.14 | 0.2 | 0.16 |
| M1: E + L | TCH | 28 | 4/24 | 0.15 | 0.13 | 0.1 | 0.11 | 0.1 | 0.16 | 0.13 |
| M1: E + L | TCH | 28 | 0/28 | 0.04 | 0.05 | 0.02 | 0 | -0.02 | 0.01 | 0.02 |
| M2: E + L + g | SA | 28 | 28/0 | 0.68 | 0.61 | 0.54 | 0.46 | 0.44 | 0.64 | 0.56 |
| M2: E + L + g | SA | 28 | 24/4 | 0.66 | 0.64 | 0.53 | 0.5 | 0.46 | 0.65 | 0.57 |
| M2: E + L + g | SA | 28 | 20/8 | 0.68 | 0.62 | 0.53 | 0.5 | 0.44 | 0.61 | 0.56 |
| M2: E + L + g | SA | 28 | 16/12 | 0.65 | 0.6 | 0.47 | 0.45 | 0.42 | 0.59 | 0.53 |
| M2: E + L + g | SA | 28 | 12/16 | 0.54 | 0.5 | 0.42 | 0.37 | 0.31 | 0.48 | 0.44 |
| M2: E + L + g | SA | 28 | 8/20 | 0.6 | 0.56 | 0.47 | 0.4 | 0.32 | 0.54 | 0.48 |
| M2: E + L + g | SA | 28 | 4/24 | 0.46 | 0.4 | 0.35 | 0.27 | 0.33 | 0.39 | 0.37 |
| M2: E + L + g | SA | 28 | 0/28 | 0.42 | 0.4 | 0.31 | 0.24 | 0.24 | 0.38 | 0.33 |
| M2: E + L + g | TCH | 28 | 28/0 | 0.34 | 0.35 | 0.21 | 0.32 | 0.29 | 0.37 | 0.31 |
| M2: E + L + g | TCH | 28 | 24/4 | 0.3 | 0.35 | 0.21 | 0.31 | 0.3 | 0.37 | 0.31 |
| M2: E + L + g | TCH | 28 | 20/8 | 0.3 | 0.35 | 0.23 | 0.3 | 0.27 | 0.35 | 0.3 |
| M2: E + L + g | TCH | 28 | 16/12 | 0.26 | 0.28 | 0.19 | 0.29 | 0.23 | 0.31 | 0.26 |
| M2: E + L + g | TCH | 28 | 12/16 | 0.23 | 0.27 | 0.14 | 0.25 | 0.16 | 0.25 | 0.22 |
| M2: E + L + g | TCH | 28 | 8/20 | 0.22 | 0.24 | 0.12 | 0.18 | 0.16 | 0.23 | 0.19 |
| M2: E + L + g | TCH | 28 | 4/24 | 0.17 | 0.14 | 0.12 | 0.12 | 0.13 | 0.19 | 0.15 |
| M2: E + L + g | TCH | 28 | 0/28 | 0.08 | 0.06 | 0.05 | 0 | 0.13 | 0.09 | 0.07 |
| M3: E + L + g + gE | SA | 28 | 28/0 | 0.72 | 0.67 | 0.56 | 0.47 | 0.44 | 0.67 | 0.59 |
| M3: E + L + g + gE | SA | 28 | 24/4 | 0.7 | 0.67 | 0.53 | 0.5 | 0.46 | 0.66 | 0.59 |
| M3: E + L + g + gE | SA | 28 | 20/8 | 0.69 | 0.64 | 0.54 | 0.5 | 0.43 | 0.62 | 0.57 |
| M3: E + L + g + gE | SA | 28 | 16/12 | 0.7 | 0.63 | 0.47 | 0.45 | 0.41 | 0.61 | 0.55 |
| M3: E + L + g + gE | SA | 28 | 12/16 | 0.57 | 0.53 | 0.4 | 0.34 | 0.32 | 0.47 | 0.44 |
| M3: E + L + g + gE | SA | 28 | 8/20 | 0.66 | 0.59 | 0.45 | 0.38 | 0.31 | 0.53 | 0.49 |
| M3: E + L + g + gE | SA | 28 | 4/24 | 0.53 | 0.47 | 0.31 | 0.22 | 0.24 | 0.41 | 0.36 |
| M3: E + L + g + gE | SA | 28 | 0/28 | 0.45 | 0.4 | 0.12 | 0.04 | 0.14 | 0.23 | 0.23 |
| M3: E + L + g + gE | TCH | 28 | 28/0 | 0.34 | 0.4 | 0.21 | 0.32 | 0.31 | 0.38 | 0.33 |
| M3: E + L + g + gE | TCH | 28 | 24/4 | 0.29 | 0.38 | 0.21 | 0.31 | 0.31 | 0.39 | 0.31 |
| M3: E + L + g + gE | TCH | 28 | 20/8 | 0.3 | 0.39 | 0.23 | 0.3 | 0.28 | 0.34 | 0.31 |
| M3: E + L + g + gE | TCH | 28 | 16/12 | 0.26 | 0.33 | 0.18 | 0.28 | 0.26 | 0.32 | 0.27 |
| M3: E + L + g + gE | TCH | 28 | 12/16 | 0.23 | 0.3 | 0.14 | 0.24 | 0.18 | 0.25 | 0.22 |
| M3: E + L + g + gE | TCH | 28 | 8/20 | 0.21 | 0.29 | 0.12 | 0.17 | 0.19 | 0.23 | 0.2 |
| M3: E + L + g + gE | TCH | 28 | 4/24 | 0.12 | 0.21 | 0.12 | 0.11 | 0.16 | 0.18 | 0.15 |
| M3: E + L + g + gE | TCH | 28 | 0/28 | 0.08 | 0.19 | 0.02 | 0.04 | 0.09 | 0.09 | 0.08 |
| M1: E + L | SA | 24 | 24/0 | 0.56 | 0.56 | 0.47 | 0.43 | 0.39 | 0.53 | 0.49 |
| M1: E + L | SA | 24 | 20/4 | 0.49 | 0.47 | 0.4 | 0.41 | 0.37 | 0.5 | 0.44 |
| M1: E + L | SA | 24 | 16/8 | 0.45 | 0.4 | 0.36 | 0.35 | 0.31 | 0.41 | 0.38 |
| M1: E + L | SA | 24 | 12/12 | 0.42 | 0.4 | 0.34 | 0.32 | 0.31 | 0.4 | 0.37 |
| M1: E + L | SA | 24 | 8/16 | 0.28 | 0.21 | 0.24 | 0.23 | 0.19 | 0.26 | 0.24 |
| M1: E + L | SA | 24 | 4/20 | 0.2 | 0.23 | 0.22 | 0.21 | 0.1 | 0.22 | 0.2 |
| M1: E + L | SA | 24 | 0/24 | -0.01 | -0.01 | -0.05 | 0.01 | -0.01 | -0.01 | -0.01 |
| M1: E + L | TCH | 24 | 24/0 | 0.28 | 0.31 | 0.17 | 0.27 | 0.28 | 0.35 | 0.28 |
| M1: E + L | TCH | 24 | 20/4 | 0.25 | 0.29 | 0.19 | 0.26 | 0.25 | 0.32 | 0.26 |
| M1: E + L | TCH | 24 | 16/8 | 0.26 | 0.27 | 0.18 | 0.23 | 0.22 | 0.3 | 0.24 |
| M1: E + L | TCH | 24 | 12/12 | 0.21 | 0.21 | 0.13 | 0.22 | 0.18 | 0.23 | 0.2 |
| M1: E + L | TCH | 24 | 8/16 | 0.15 | 0.19 | 0.09 | 0.19 | 0.12 | 0.19 | 0.15 |
| M1: E + L | TCH | 24 | 4/20 | 0.13 | 0.1 | 0.08 | 0.13 | 0.08 | 0.11 | 0.1 |
| M1: E + L | TCH | 24 | 0/24 | -0.04 | -0.03 | -0.01 | 0 | 0.01 | 0 | -0.01 |
| M2: E + L + g | SA | 24 | 24/0 | 0.67 | 0.65 | 0.54 | 0.48 | 0.44 | 0.61 | 0.57 |
| M2: E + L + g | SA | 24 | 20/4 | 0.64 | 0.6 | 0.49 | 0.47 | 0.43 | 0.61 | 0.54 |
| M2: E + L + g | SA | 24 | 16/8 | 0.63 | 0.57 | 0.47 | 0.43 | 0.39 | 0.56 | 0.51 |
| M2: E + L + g | SA | 24 | 12/12 | 0.62 | 0.57 | 0.46 | 0.41 | 0.39 | 0.56 | 0.5 |
| M2: E + L + g | SA | 24 | 8/16 | 0.47 | 0.42 | 0.37 | 0.32 | 0.29 | 0.42 | 0.38 |
| M2: E + L + g | SA | 24 | 4/20 | 0.44 | 0.42 | 0.36 | 0.3 | 0.24 | 0.4 | 0.36 |
| M2: E + L + g | SA | 24 | 0/24 | 0.41 | 0.36 | 0.29 | 0.2 | 0.27 | 0.34 | 0.31 |
| M2: E + L + g | TCH | 24 | 24/0 | 0.3 | 0.33 | 0.18 | 0.27 | 0.29 | 0.37 | 0.29 |
| M2: E + L + g | TCH | 24 | 20/4 | 0.27 | 0.32 | 0.21 | 0.28 | 0.27 | 0.34 | 0.28 |
| M2: E + L + g | TCH | 24 | 16/8 | 0.28 | 0.3 | 0.2 | 0.25 | 0.23 | 0.31 | 0.26 |
| M2: E + L + g | TCH | 24 | 12/12 | 0.23 | 0.24 | 0.14 | 0.23 | 0.19 | 0.25 | 0.21 |
| M2: E + L + g | TCH | 24 | 8/16 | 0.18 | 0.24 | 0.12 | 0.22 | 0.13 | 0.22 | 0.18 |
| M2: E + L + g | TCH | 24 | 4/20 | 0.18 | 0.17 | 0.1 | 0.17 | 0.1 | 0.16 | 0.15 |
| M2: E + L + g | TCH | 24 | 0/24 | 0.07 | 0.11 | 0.06 | 0.06 | 0.06 | 0.09 | 0.07 |
| M3: E + L + g + gE | SA | 24 | 24/0 | 0.72 | 0.68 | 0.56 | 0.49 | 0.45 | 0.65 | 0.59 |
| M3: E + L + g + gE | SA | 24 | 20/4 | 0.66 | 0.63 | 0.5 | 0.46 | 0.44 | 0.63 | 0.55 |
| M3: E + L + g + gE | SA | 24 | 16/8 | 0.65 | 0.6 | 0.47 | 0.42 | 0.38 | 0.55 | 0.51 |
| M3: E + L + g + gE | SA | 24 | 12/12 | 0.62 | 0.56 | 0.44 | 0.4 | 0.37 | 0.56 | 0.49 |
| M3: E + L + g + gE | SA | 24 | 8/16 | 0.52 | 0.47 | 0.31 | 0.28 | 0.28 | 0.4 | 0.38 |
| M3: E + L + g + gE | SA | 24 | 4/20 | 0.5 | 0.47 | 0.3 | 0.25 | 0.2 | 0.4 | 0.35 |
| M3: E + L + g + gE | SA | 24 | 0/24 | 0.45 | 0.42 | 0.15 | 0.05 | 0.04 | 0.2 | 0.22 |
| M3: E + L + g + gE | TCH | 24 | 24/0 | 0.31 | 0.35 | 0.19 | 0.27 | 0.31 | 0.38 | 0.3 |
| M3: E + L + g + gE | TCH | 24 | 20/4 | 0.27 | 0.37 | 0.2 | 0.26 | 0.28 | 0.36 | 0.29 |
| M3: E + L + g + gE | TCH | 24 | 16/8 | 0.26 | 0.32 | 0.18 | 0.24 | 0.26 | 0.32 | 0.26 |
| M3: E + L + g + gE | TCH | 24 | 12/12 | 0.2 | 0.27 | 0.14 | 0.23 | 0.22 | 0.26 | 0.22 |
| M3: E + L + g + gE | TCH | 24 | 8/16 | 0.13 | 0.29 | 0.11 | 0.19 | 0.16 | 0.21 | 0.18 |
| M3: E + L + g + gE | TCH | 24 | 4/20 | 0.15 | 0.24 | 0.09 | 0.13 | 0.14 | 0.16 | 0.15 |
| M3: E + L + g + gE | TCH | 24 | 0/24 | 0.03 | 0.18 | 0.06 | -0.02 | 0.1 | 0.08 | 0.07 |
| M1: E + L | SA | 20 | 20/0 | 0.45 | 0.49 | 0.43 | 0.38 | 0.34 | 0.49 | 0.43 |
| M1: E + L | SA | 20 | 16/4 | 0.42 | 0.4 | 0.35 | 0.35 | 0.35 | 0.45 | 0.39 |
| M1: E + L | SA | 20 | 12/8 | 0.39 | 0.33 | 0.31 | 0.31 | 0.26 | 0.36 | 0.33 |
| M1: E + L | SA | 20 | 8/12 | 0.35 | 0.35 | 0.3 | 0.3 | 0.26 | 0.36 | 0.32 |
| M1: E + L | SA | 20 | 4/16 | 0.24 | 0.16 | 0.2 | 0.19 | 0.14 | 0.23 | 0.19 |
| M1: E + L | SA | 20 | 0/20 | -0.03 | -0.01 | -0.02 | 0.01 | -0.02 | -0.02 | -0.01 |
| M1: E + L | TCH | 20 | 20/0 | 0.26 | 0.29 | 0.14 | 0.27 | 0.27 | 0.32 | 0.26 |
| M1: E + L | TCH | 20 | 16/4 | 0.21 | 0.26 | 0.18 | 0.24 | 0.23 | 0.27 | 0.23 |
| M1: E + L | TCH | 20 | 12/8 | 0.19 | 0.25 | 0.17 | 0.21 | 0.18 | 0.25 | 0.21 |
| M1: E + L | TCH | 20 | 8/12 | 0.18 | 0.18 | 0.11 | 0.18 | 0.14 | 0.19 | 0.16 |
| M1: E + L | TCH | 20 | 4/16 | 0.11 | 0.12 | 0.08 | 0.11 | 0.07 | 0.14 | 0.1 |
| M1: E + L | TCH | 20 | 0/20 | -0.01 | 0.03 | -0.02 | 0 | 0.02 | 0.04 | 0.01 |
| M2: E + L + g | SA | 20 | 20/0 | 0.59 | 0.59 | 0.5 | 0.44 | 0.39 | 0.59 | 0.52 |
| M2: E + L + g | SA | 20 | 16/4 | 0.58 | 0.55 | 0.45 | 0.42 | 0.42 | 0.57 | 0.5 |
| M2: E + L + g | SA | 20 | 12/8 | 0.56 | 0.5 | 0.42 | 0.39 | 0.34 | 0.5 | 0.45 |
| M2: E + L + g | SA | 20 | 8/12 | 0.58 | 0.54 | 0.44 | 0.4 | 0.35 | 0.54 | 0.47 |
| M2: E + L + g | SA | 20 | 4/16 | 0.41 | 0.35 | 0.31 | 0.26 | 0.23 | 0.36 | 0.32 |
| M2: E + L + g | SA | 20 | 0/20 | 0.25 | 0.23 | 0.21 | 0.13 | 0.18 | 0.22 | 0.2 |
| M2: E + L + g | TCH | 20 | 20/0 | 0.28 | 0.31 | 0.15 | 0.28 | 0.28 | 0.33 | 0.27 |
| M2: E + L + g | TCH | 20 | 16/4 | 0.24 | 0.28 | 0.19 | 0.25 | 0.25 | 0.29 | 0.25 |
| M2: E + L + g | TCH | 20 | 12/8 | 0.22 | 0.27 | 0.18 | 0.23 | 0.2 | 0.27 | 0.23 |
| M2: E + L + g | TCH | 20 | 8/12 | 0.2 | 0.22 | 0.14 | 0.2 | 0.15 | 0.22 | 0.19 |
| M2: E + L + g | TCH | 20 | 4/16 | 0.15 | 0.16 | 0.1 | 0.13 | 0.08 | 0.16 | 0.13 |
| M2: E + L + g | TCH | 20 | 0/20 | 0.03 | 0.06 | 0.03 | 0.05 | 0.08 | 0.07 | 0.05 |
| M3: E + L + g + gE | SA | 20 | 20/0 | 0.68 | 0.59 | 0.52 | 0.43 | 0.39 | 0.6 | 0.54 |
| M3: E + L + g + gE | SA | 20 | 16/4 | 0.6 | 0.59 | 0.43 | 0.41 | 0.41 | 0.57 | 0.5 |
| M3: E + L + g + gE | SA | 20 | 12/8 | 0.61 | 0.52 | 0.41 | 0.34 | 0.31 | 0.48 | 0.44 |
| M3: E + L + g + gE | SA | 20 | 8/12 | 0.64 | 0.57 | 0.4 | 0.37 | 0.34 | 0.56 | 0.48 |
| M3: E + L + g + gE | SA | 20 | 4/16 | 0.45 | 0.42 | 0.25 | 0.22 | 0.22 | 0.35 | 0.32 |
| M3: E + L + g + gE | SA | 20 | 0/20 | 0.33 | 0.26 | -0.06 | -0.03 | 0.13 | 0.11 | 0.13 |
| M3: E + L + g + gE | TCH | 20 | 20/0 | 0.28 | 0.33 | 0.15 | 0.28 | 0.3 | 0.34 | 0.28 |
| M3: E + L + g + gE | TCH | 20 | 16/4 | 0.22 | 0.33 | 0.19 | 0.25 | 0.26 | 0.31 | 0.26 |
| M3: E + L + g + gE | TCH | 20 | 12/8 | 0.19 | 0.3 | 0.19 | 0.23 | 0.22 | 0.26 | 0.23 |
| M3: E + L + g + gE | TCH | 20 | 8/12 | 0.19 | 0.26 | 0.12 | 0.19 | 0.18 | 0.23 | 0.19 |
| M3: E + L + g + gE | TCH | 20 | 4/16 | 0.11 | 0.22 | 0.07 | 0.1 | 0.13 | 0.15 | 0.13 |
| M3: E + L + g + gE | TCH | 20 | 0/20 | 0.04 | 0.13 | 0.05 | 0.02 | 0.1 | 0.04 | 0.06 |
| M1: E + L | SA | 16 | 16/0 | 0.41 | 0.41 | 0.36 | 0.31 | 0.32 | 0.36 | 0.36 |
| M1: E + L | SA | 16 | 12/4 | 0.34 | 0.31 | 0.31 | 0.3 | 0.29 | 0.38 | 0.32 |
| M1: E + L | SA | 16 | 8/8 | 0.29 | 0.27 | 0.24 | 0.24 | 0.21 | 0.27 | 0.25 |
| M1: E + L | SA | 16 | 4/12 | 0.19 | 0.2 | 0.17 | 0.18 | 0.19 | 0.25 | 0.2 |
| M1: E + L | SA | 16 | 0/16 | -0.01 | -0.02 | -0.03 | -0.04 | -0.01 | -0.02 | -0.02 |
| M1: E + L | TCH | 16 | 16/0 | 0.22 | 0.23 | 0.11 | 0.2 | 0.17 | 0.23 | 0.19 |
| M1: E + L | TCH | 16 | 12/4 | 0.18 | 0.2 | 0.16 | 0.22 | 0.18 | 0.24 | 0.2 |
| M1: E + L | TCH | 16 | 8/8 | 0.17 | 0.21 | 0.12 | 0.15 | 0.16 | 0.2 | 0.17 |
| M1: E + L | TCH | 16 | 4/12 | 0.11 | 0.1 | 0.05 | 0.09 | 0.08 | 0.13 | 0.09 |
| M1: E + L | TCH | 16 | 0/16 | -0.07 | -0.05 | -0.02 | -0.02 | -0.01 | -0.03 | -0.03 |
| M2: E + L + g | SA | 16 | 16/0 | 0.55 | 0.52 | 0.44 | 0.39 | 0.37 | 0.47 | 0.46 |
| M2: E + L + g | SA | 16 | 12/4 | 0.51 | 0.47 | 0.4 | 0.37 | 0.36 | 0.5 | 0.44 |
| M2: E + L + g | SA | 16 | 8/8 | 0.48 | 0.45 | 0.36 | 0.33 | 0.3 | 0.43 | 0.39 |
| M2: E + L + g | SA | 16 | 4/12 | 0.37 | 0.34 | 0.28 | 0.26 | 0.27 | 0.38 | 0.32 |
| M2: E + L + g | SA | 16 | 0/16 | 0.31 | 0.29 | 0.23 | 0.17 | 0.21 | 0.28 | 0.25 |
| M2: E + L + g | TCH | 16 | 16/0 | 0.23 | 0.25 | 0.12 | 0.21 | 0.19 | 0.25 | 0.21 |
| M2: E + L + g | TCH | 16 | 12/4 | 0.2 | 0.23 | 0.18 | 0.24 | 0.2 | 0.26 | 0.22 |
| M2: E + L + g | TCH | 16 | 8/8 | 0.19 | 0.23 | 0.14 | 0.17 | 0.18 | 0.22 | 0.19 |
| M2: E + L + g | TCH | 16 | 4/12 | 0.13 | 0.14 | 0.07 | 0.11 | 0.09 | 0.15 | 0.12 |
| M2: E + L + g | TCH | 16 | 0/16 | 0.03 | 0.07 | 0.05 | 0.05 | 0.01 | 0.07 | 0.05 |
| M3: E + L + g + gE | SA | 16 | 16/0 | 0.61 | 0.51 | 0.43 | 0.35 | 0.38 | 0.48 | 0.46 |
| M3: E + L + g + gE | SA | 16 | 12/4 | 0.54 | 0.54 | 0.38 | 0.32 | 0.34 | 0.46 | 0.43 |
| M3: E + L + g + gE | SA | 16 | 8/8 | 0.53 | 0.48 | 0.35 | 0.28 | 0.22 | 0.43 | 0.38 |
| M3: E + L + g + gE | SA | 16 | 4/12 | 0.47 | 0.39 | 0.25 | 0.21 | 0.26 | 0.35 | 0.32 |
| M3: E + L + g + gE | SA | 16 | 0/16 | 0.32 | 0.25 | -0.05 | -0.03 | 0.12 | 0.17 | 0.13 |
| M3: E + L + g + gE | TCH | 16 | 16/0 | 0.21 | 0.27 | 0.13 | 0.21 | 0.21 | 0.25 | 0.21 |
| M3: E + L + g + gE | TCH | 16 | 12/4 | 0.18 | 0.26 | 0.18 | 0.23 | 0.2 | 0.26 | 0.22 |
| M3: E + L + g + gE | TCH | 16 | 8/8 | 0.18 | 0.25 | 0.14 | 0.16 | 0.21 | 0.21 | 0.19 |
| M3: E + L + g + gE | TCH | 16 | 4/12 | 0.09 | 0.19 | 0.05 | 0.08 | 0.14 | 0.16 | 0.12 |
| M3: E + L + g + gE | TCH | 16 | 0/16 | 0.05 | 0.12 | 0.03 | 0 | 0.08 | 0.07 | 0.06 |
| M1: E + L | SA | 12 | 12/0 | 0.47 | 0.4 | 0.37 | 0.37 | 0.31 | 0.44 | 0.39 |
| M1: E + L | SA | 12 | 8/4 | 0.26 | 0.23 | 0.22 | 0.21 | 0.23 | 0.26 | 0.23 |
| M1: E + L | SA | 12 | 4/8 | 0.2 | 0.18 | 0.16 | 0.17 | 0.13 | 0.2 | 0.17 |
| M1: E + L | SA | 12 | 0/12 | 0.03 | 0.02 | -0.01 | 0.01 | 0.03 | 0.05 | 0.02 |
| M1: E + L | TCH | 12 | 12/0 | 0.22 | 0.23 | 0.14 | 0.23 | 0.19 | 0.22 | 0.2 |
| M1: E + L | TCH | 12 | 8/4 | 0.13 | 0.17 | 0.15 | 0.16 | 0.18 | 0.23 | 0.17 |
| M1: E + L | TCH | 12 | 4/8 | 0.12 | 0.14 | 0.09 | 0.13 | 0.11 | 0.12 | 0.12 |
| M1: E + L | TCH | 12 | 0/12 | 0 | 0 | 0.01 | 0 | -0.02 | -0.02 | 0 |
| M2: E + L + g | SA | 12 | 12/0 | 0.63 | 0.56 | 0.48 | 0.44 | 0.38 | 0.58 | 0.51 |
| M2: E + L + g | SA | 12 | 8/4 | 0.36 | 0.34 | 0.29 | 0.25 | 0.27 | 0.34 | 0.31 |
| M2: E + L + g | SA | 12 | 4/8 | 0.34 | 0.31 | 0.25 | 0.24 | 0.2 | 0.31 | 0.27 |
| M2: E + L + g | SA | 12 | 0/12 | 0.29 | 0.25 | 0.21 | 0.14 | 0.12 | 0.2 | 0.2 |
| M2: E + L + g | TCH | 12 | 12/0 | 0.23 | 0.25 | 0.16 | 0.24 | 0.2 | 0.24 | 0.22 |
| M2: E + L + g | TCH | 12 | 8/4 | 0.15 | 0.19 | 0.16 | 0.18 | 0.19 | 0.25 | 0.19 |
| M2: E + L + g | TCH | 12 | 4/8 | 0.15 | 0.17 | 0.1 | 0.15 | 0.13 | 0.15 | 0.14 |
| M2: E + L + g | TCH | 12 | 0/12 | 0.02 | 0.08 | 0.03 | 0.02 | 0.03 | 0.06 | 0.04 |
| M3: E + L + g + gE | SA | 12 | 12/0 | 0.64 | 0.58 | 0.49 | 0.43 | 0.38 | 0.55 | 0.51 |
| M3: E + L + g + gE | SA | 12 | 8/4 | 0.39 | 0.37 | 0.29 | 0.2 | 0.26 | 0.32 | 0.31 |
| M3: E + L + g + gE | SA | 12 | 4/8 | 0.39 | 0.33 | 0.23 | 0.18 | 0.14 | 0.28 | 0.26 |
| M3: E + L + g + gE | SA | 12 | 0/12 | 0.34 | 0.31 | 0.03 | -0.04 | 0.11 | 0.12 | 0.14 |
| M3: E + L + g + gE | TCH | 12 | 12/0 | 0.22 | 0.29 | 0.16 | 0.25 | 0.24 | 0.26 | 0.24 |
| M3: E + L + g + gE | TCH | 12 | 8/4 | 0.15 | 0.22 | 0.16 | 0.18 | 0.21 | 0.26 | 0.2 |
| M3: E + L + g + gE | TCH | 12 | 4/8 | 0.14 | 0.2 | 0.09 | 0.13 | 0.16 | 0.14 | 0.14 |
| M3: E + L + g + gE | TCH | 12 | 0/12 | 0.06 | 0.16 | 0 | 0.01 | 0.09 | 0.05 | 0.06 |

**Supplementary Table 2.** Statistical significancy of model’s performance differences. The following ANOVA test was run for i) all results and ii) only the results with the maximum calibration size (186 phenotypes in the train set): accuracy = model + environment + design. Tukey’s HSD was then applied to the ANOVA object to test differences between models. No significant differences between models M2 and M3 were found except for TCH and maximum calibration size.

| Dataset | trait | contrast | diff | lwr | upr | p.adj |
| --- | --- | --- | --- | --- | --- | --- |
| all<br>calibration<br>sizes | SA | M3-M2 | -0.001 | -0.023 | 0.02 | 0.988 |
|  |  | M1-M2 | -0.135 | -0.157 | -0.113 | 0 |
|  |  | M1-M3 | -0.134 | -0.155 | -0.112 | 0 |
|  | TCH | M3-M2 | 0.007 | -0.004 | 0.018 | 0.293 |
|  |  | M1-M2 | -0.023 | -0.034 | -0.011 | 0 |
|  |  | M1-M3 | -0.03 | -0.041 | -0.019 | 0 |
| max<br>calibration<br>size (186) | SA | M3-M2 | -0.01 | -0.022 | 0.001 | 0.094 |
|  |  | M1-M2 | -0.14 | -0.152 | -0.129 | 0 |
|  |  | M1-M3 | -0.13 | -0.142 | -0.118 | 0 |
|  | TCH | M3-M2 | 0.007 | 0.002 | 0.013 | 0.008 |
|  |  | M1-M2 | -0.026 | -0.032 | -0.021 | 0 |
|  |  | M1-M3 | -0.033 | -0.039 | -0.028 | 0 |

